# A distributed algorithm to maintain and repair the trail networks of arboreal ants

**DOI:** 10.1101/194480

**Authors:** Arjun Chandrasekhar, Deborah M. Gordon, Saket Navlakha

## Abstract

We study how the arboreal turtle ant (*Cephalotes goniodontus*) solves a fundamental computing problem: maintaining a trail network and finding alternative paths to route around broken links in the network. Turtle ants form a routing backbone of foraging trails linking several nests and temporary food sources. This species travels only in the trees, so their foraging trails are constrained to lie on a natural graph formed by overlapping branches and vines in the tangled canopy. Links between branches, however, can be ephemeral, easily destroyed by wind, rain, or animal movements. Here we report a biologically feasible distributed algorithm, parameterized using field data, that can plausibly describe how turtle ants maintain the routing backbone and find alternative paths to circumvent broken links in the backbone. We validate the ability of this probabilistic algorithm to circumvent simulated breaks in synthetic and real-world networks, and we derive an analytic explanation for why certain features are crucial to improve the algorithm’s success. Our proposed algorithm uses fewer computational resources than common distributed graph search algorithms, and thus may be useful in other domains, such as for swarm computing or for coordinating molecular robots.

## Introduction

Distributed algorithms allow a collection of agents to efficiently solve computational problems without centralized control [1]. Recent research has uncovered such algorithms implemented by many biological systems, including slime molds during foraging [2] and neural circuits during development [3]. Ants are a diverse taxon of more than 14,000 species that have also evolved distributed algorithms to establish trail networks [4]. Investigating the algorithms used by biological systems can reveal novel solutions to engineering problems [5, 3].

Here we present the first computational analysis, parameterized using data from field observations, of trail networks of an arboreal ant species. The arboreal turtle ant *C. goniodontus* nests and forages in the trees in the tropical dry forest of western Mexico [6]. Because the ants never leave the trees, their foraging trails are constrained by a natural graph: branches and vines form the edges in the graph, and junctions at overlapping branches form the nodes (Figure 1A–C). Each colony has several nests, located in dead tree branches, that are connected to each other in a circuit or network routing backbone [4, 7, 8]. Moving on the trails along this backbone, the ants distribute resources among the juveniles, workers, and reproductives in all of the nests, while additional temporary trails split from the backbone and lead to food sources. The backbone trail network can be large, often extending over 50 meters in circumference, and encompassing numerous trees [6]. The ants use many junctions in dense vegetation, so trails can be tortuous; each meter of linear distance typically requires ants to traverse approximately 2–5 meters of vegetation [6]. The colony thus chooses paths in the network from a myriad of potential routes, dictated by the graph structure of the vegetation. Ants lay trail pheromone as they move along the edges, and ants use pheromone when choosing edges.

We present a distributed algorithm that can plausibly describe how turtle ants maintain and repair breaks to their routing backbone. Links between branches or vines can be ephemeral, often disrupted by wind, rain, or the movement of an animal through the vegetation. To re-establish connectivity of the routing backbone after a break, the ants must establish a new path that reconnects the two sides of the broken trail. This is an important problem in many network applications [9] and can be solved efficiently using numerous graph algorithms, such as Dijkstra’s algorithm or the Bellman-Ford algorithm [10]. However, these classic algorithms require significantly more computation and memory than is likely available to simple biological agents such as turtle ants, who regulate their behavior using local interactions rather than central control [11].

Repairing breaks requires overcoming three challenges. First, the ants must succeed in finding an alternative path by exploring new edges that currently have no pheromone and avoiding deadends in the network. One hypothesis for how this could be achieved is to first generate many candidate alternative paths and then converge to one or a few of them over time — a process we call “pruning”. Such a strategy, also employed by slime molds [2] and neural circuits [3], has been shown to help quickly discover new paths in distributed settings, in which no agent is aware of the topology of the entire network. Second, all ants must converge to the same new path in order to optimally coordinate resource transport. Turtle ants travel in coherent trails that link nests and food sources [12]. After the vegetation supporting the trail is ruptured, the ants explore outside the previous path, and eventually commit again to a single path. Such convergence prevents ants from getting lost or separated from the rest of the colony. This is also an important goal in computer routing networks, where convergence to a single path ensures in-order delivery of data packets [13]. Third, it may be important to minimize the length of the new trail, which is also a standard measure of efficiency used when evaluating transport network design. However, data from field studies [6, 12] suggest that turtle ant paths are often not the shortest globally. It appears that the second objective, successful convergence, is more important than minimizing trail length, presumably because ants getting lost or separated has a higher cost than the energy spent in walking [12]. A common strategy to increase robustness to edge failures in a graph is to include loops in the path. Prior work [12] showed that loops do form in turtle ant trail networks; however, loops tend to get pruned over time, perhaps reducing the number of foragers needed to maintain the path.

The distributed algorithm used to maintain and repair trail networks must be robust across varying planar network topologies. The forest canopy is highly complex and dynamic, and it is unlikely that turtle ants use different algorithms to accommodate different network structures. Thus, we seek an algorithm that, while likely not “optimal” for any single planar topology, performs well across different planar topologies. The algorithm must also use very limited memory of individual agents, as ants are not capable of remembering many of their steps along the graph structure.

Our work seeks to uncover a biologically plausible distributed algorithm that corresponds with field observations of turtle ant behavior in response to experimentally-induced edge breaks (Figure 1B) [12]. We ask:

1. What model is most likely to explain how turtle ants at a node select which edge to traverse next?
2. How well can the algorithm repair broken trails in simulated breaks in synthetic and realworld network topologies when parameterized by the most biologically realistic parameter values?
  (a) Does the algorithm consistently converge to a single consensus path?
  (b) Does the algorithm find short paths?
  (c) Does bi-directional search, using ants from both sides of the broken path concurrently, improve the performance of the algorithm relative to uni-directional search?
  (d) How does allowing an ant to avoid going back to the node it previously visited (backtracking), improve algorithm performance relative to performance when ants are not prevented from backtracking?
  (e) Can we provide any theoretical insights into why certain model features are necessary for any plausible turtle ant algorithm?
3. Can the same algorithm used to repair breaks also be used to keep the established routing backbone intact in the absence of a break?
4. Do turtle ants form multiple alternative paths and then prune some of them over time, as also observed in field studies?

A model that performs well on all of these criteria can be considered a plausible model of turtle ant behavior. Our main contribution is to identify several plausible non-linear models; we also show why one common linear model is likely implausible despite succeeding on some of the criteria listed above.

## Related work

To our knowledge, this is the first computational analysis of trail networks of an arboreal ant species, whose movements are constrained to a discrete graph structure rather than continuous space. Compared to previous work, we attempt to solve the network repair problem using different constraints and fewer assumptions about the computational abilities of individual ants.

### Species-specific modeling of ant behavior

Previous studies of ant trail networks have largely examined species that forage on a continuous 2D surface [14], including Pharaoh’s ants [15], Argentine ants [4, 16, 17], leaf-cutter ants [18], army ants [19], and red wood ants [20]. These species can define nodes and edges at any location on the surface, and form trails using techniques such as random amplification [21, 22, 19], or using their own bodies to form living bridges [23]. Experimental work on these species sometimes uses discrete mazes or Y-junctions to impose a graph structure; however, these species have evolved to create graph structures in continuous space, not to solve problems on a fixed graph structure, as turtle ants have evolved to do. Turtle ant movements are entirely constrained by the vegetation in which they travel. They cannot form trails with nodes and edges at arbitrary locations; instead, they can use only the nodes and edges that are available to them.

Further, to provide the simplest possible algorithm that is biologically realistic, we assume that turtle ants use only one type of pheromone. There are more than 14,000 species of ants, and they differ in their use of chemical cues. For example, *Monomorium pharoensis* uses several different trail pheromones [24, 25, 26, 27, 28]. There is, however, no evidence that turtle ants lay more than one type of trail pheromone.

### Ant colony optimization

Models of ant colony optimization (ACO), first proposed in 1991, loosely mimic ant behavior to solve combinatorial optimization problems, such as the traveling salesman problem [29, 30, 31]. In ACO, individual ants each use a heuristic to construct candidate solutions, and then use pheromone to lead other ants towards higher quality solutions. Recent advances improve ACO through techniques such as local search [32], cunning ants [33], and iterated ants [34]. ACO, however, provides simulated ants more computational power than turtle actually ants possess; in particular, ACO-simulated ants have sufficient memory to remember, retrace, and reinforce entire paths or solutions, and they can choose how much pheromone to lay in retrospect, based on the optimality of the solution.

Prior work inspired by ants provides solutions to graph search problems [35, 36], such as the Hamiltonian path problem [37] or the Ants Nearby Treasure Search (ANTS) problem. The latter investigates how simulated ants collaboratively search the integer plane for a treasure source. These models afford the simulated ants various computational abilities, including searching exhaustively around a fixed radius [38], sending constant sized messages [39], or laying pheromone to mark an edge as explored [40]. Our work involves a similar model of distributed computation, but our problem requires not only that the ants find an alternative path to a nest (a “treasure”), but also that all the ants commit to using the same alternative path. This requires a fundamentally different strategy from that required for just one ant to find a treasure.

### Graph algorithms and reinforced random walks

Common algorithms used to solve the general network search and repair problem, including Dijkstra’s algorithm, breadth-first search, depth-first search, and A* search [10], all require substantial communication or memory complexity. For example, agents must maintain a large routing table, store and query a list of all previously visited nodes, or pre-compute a topology-dependent heuristic to compute node-to-node distances [41]. These abilities are all unlikely for turtle ants.

Distributed graph algorithms, in which nodes are treated as fixed agents capable of passing messages to neighbors, have also been proposed to find shortest paths in a graph [42, 43], to construct minimum spanning trees [44, 45], and to approximate various NP-hard problems [46, 47]. In contrast, our work uses a more restrictive model of distributed computation, where agents communicate only through pheromone which does not have a specific targeted recipient.

Finally, the limited assumptions about the memory of turtle ants invite comparison to a Markov process. Edge-reinforced random walks [48], first introduced by Diaconis and others [49, 50], proceed as follows: an agent, or random walker, traverses a graph by choosing amongst adjacent edges with a probability proportional to their edge weight; then the agent augments the weight (or pheromone) of each edge chosen. Our model expands edge-reinforced random walks in two ways: first, we allow many agents to walk the graph concurrently, and second, we decrease edge weights over time. Our work is similar to previous models of the gliding behavior of myxobacteria [51] that consider synchronous, node (rather than edge)-reinforced, random walks with decay. These models seek to determine when bacteria aggregate on adjacent points or instead walk freely on the grid. By contrast, here we ask whether the random walkers converge to a single consensus path between two points on the grid that are not necessarily adjacent.

## Results

Our goals are to find an algorithm that can simultaneously explain the movement patterns of turtle ants on a trail network and that can effectively solve the network repair problem. First, we describe a computational framework for evaluating the collective response of turtle ants to edge ruptures. We evaluate the response according to three objectives: the likelihood of finding an alternative path to repair the trail, how well the ants converge to the same new trail, and the capacity to minimize the length of the trail. Second, we derive multiple candidate distributed algorithms for network repair. We parameterize each algorithm using data from field experiments to determine how the model would predict which edge a turtle ant would choose to traverse next from a node, given only local information about adjacent edges and their edge weights. Third, we analyze via simulation how our algorithms perform on different planar network topologies, including simulated breaks on a European road transport network.

### A graph-theoretic framework for modeling network repair by turtle ants

We start with a weighted, undirected graph *G* = (*V, E, W*), where *V* is the node set, *E* is the edge set, and *W* are the edge weights, as well as two nest nodes *u, v ∈ V*, and a path *P* = (*u, …, v*) from *u* to *v* with pheromone along each edge in *P*. Edges are undirected since turtle ants can walk in both directions over edges. Edge weights correspond to the amount of pheromone on the edges, which can change over time. We mimic a break in the path by removing some edge in *P*. The challenge for the ants is to find an alternative path that reconnects *u* and *v*. The alternative path may build off the existing path, so that the initial and final path may share some edges.

Communication among simulated ants is limited to chemical signals, analogous to pheromone, left on edges traversed. Field observations are consistent with the assumption that, like Argentine ants [52], turtle ants lay trail pheromone continuously as they walk [6, 12]. Though this has not been observed directly, we hypothesize that there are certain exceptional situations in which turtle ants discontinue laying pheromone (Methods). Each ant at a node senses the pheromone level on adjacent edges to inform its next movement. Observations suggest that a turtle ant tends to keep moving in the same direction, indicating that an ant is able to avoid the previous node it visited. We thus assume that simulated ants have one time-step of memory, used to avoid going back and forth along the same edge. As is characteristic of many species of ants [53, 54, 55], simulated ants have no unique identifiers and can use only local information.

#### Parameters

Our algorithm uses three biological parameters: *q*_add_, *q*_decay_, and *q*_explore_.

The first parameter (*q*_add_) determines how much pheromone is added when an ant traverses an edge. After each time step, each edge (*v*_1_, *v*_2_) traversed increases its edge weight as:

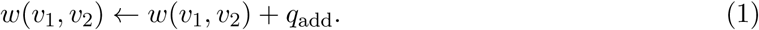

Without loss of generality, we fix *q*_add_ = 1, representing a unit of pheromone that an ant deposits on each edge traversed.

The second parameter (*q*_decay_) specifies how much pheromone evaporates on each edge in each time step due to natural decay. We model pheromone decay as an exponential decrease in edge weight [56, 57]; thus *q*_decay_ ∈ (0, 1), and at each time step, for each edge (*v*_1_, *v*_2_), its weight is updated as:

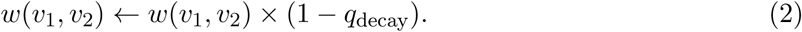

Larger values of *q*_decay_ correspond to more rapid decay of pheromone on the edge.

The third parameter (*q*_explore_) specifies the probability that an ant takes an “explore step”. The definition of an “explore step” is algorithm-specific (see below), but intuitively, it involves choosing an edge with relatively less or no pheromone. Such deviation is clearly required by any network repair algorithm, since after the routing backbone is ruptured, edges not part of the existing path must be traversed to repair the break. Field observations show that even in the absence of a break, turtle ants explore edges off the main trail. This allows them to discover new food sources and incorporate them into the trail network [12].

#### Performance metrics

After *T* time steps, we evaluate the outcome of the algorithm using the following measures (averaged over 50 repeat simulations):

1. <u>Success rate</u>: The probability that the simulated ants succeeded in forming a new path from *u* to *v* that does not use the broken edge. In this new path, ants are not required to traverse edges of relatively low weight (Methods). Higher values are better; for example, a success rate of 70% means that in 70% of the simulations, the ants successfully formed an alternative path.
2. <u>Path entropy</u>: An information-theoretic measure of how well the ants converge to a single consensus path, rather than creating multiple, potentially overlapping, *u → v* paths with pheromone. Lower values are better, indicating that subsequent ants using the same algorithm on the resulting network will all follow a common path, rather than dispersing along many different paths. This measure is computed only in the simulations in which an alternative path was successfully found. Field observations show that turtle ants consistently converge to a consensus path, and loops in the network are often pruned away over time [12]. This reduces the numbers of lost ants and the numbers of ants traveling in circles.
3. <u>Path length</u>: The length of the new path. Although turtle ants do not always find the globally shortest path [12], we include this measure because it is commonly used to evaluate routing algorithms. Lower values are better, indicating shorter paths. This measure is computed only in the simulations in which an alternative path was successfully found.

#### A model of computation for individual ants

We assume that all ants are identical and have the following computational abilities:

- Each ant can avoid the node it immediately previously visited. It cannot, however, remember its entire path from the nest up to its current point. The ant may also keep track of a binary state variable that determines whether it is combing back from a dead end and should discontinue laying pheromone.
- In field observations, ants appear to pause at nodes and inspect more than one edge before choosing an edge to take [12]. Thus, each ant can access all adjacent edge weights to decide which node to visit next. To choose its next edge, we allow ants to perform any Turing-computable computation, although we show that a simple, albeit non-linear, function will suffice.

See Methods for full technical details of the model and performance metrics.

### Candidate distributed algorithms

Below we introduce several biologically plausible algorithms that attempt to describe how a turtle ant at a node *s* chooses which edge to traverse next among possible neighboring edges *t*_1_, *t*_2_, *…t*_*n*_. These algorithms build upon previous linear and non-linear models used to analyze ant trail formation in other species, such as Argentine ants [58, 59, 60] and pharaoh ants [61]. Let *w*(*s, t*_*i*_) be the current weight on edge (*s, t*_*i*_), and let uniform() be a random value drawn uniformly from [0, 1].

In the WEIGHTED random walk (Algorithm 1), each ant chooses the next edge to traverse with probability proportional to the amount of pheromone on that edge: the more pheromone on an edge, the more likely an ant is to traverse that edge. However, with probability *q*_explore_, the ant takes an edge that has zero pheromone.

**Algorithm 1.**
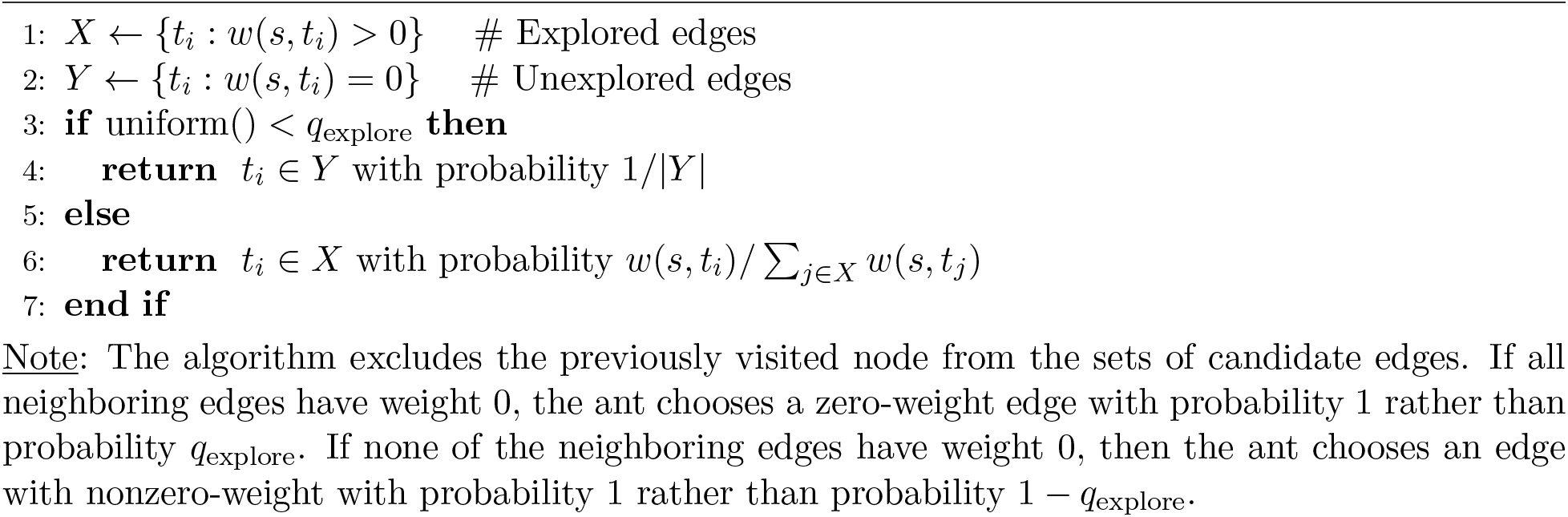
WEIGHTED random walk.

In the RANKEDGE random walk (Algorithm 2), with probability 1 *− q*_explore_, the ant chooses an edge with the highest weight (ties are broken at random). With probability *q*_explore_, it bypasses the highest weighted edges and considers edges with the second highest weight. With probability *q*_explore_(1 *− q*_explore_), it chooses an edge with the second highest weight. With probability 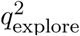, it bypasses both the highest and second highest weighted edges and considers edges with the third highest weight, and so on.

**Algorithm 2.**
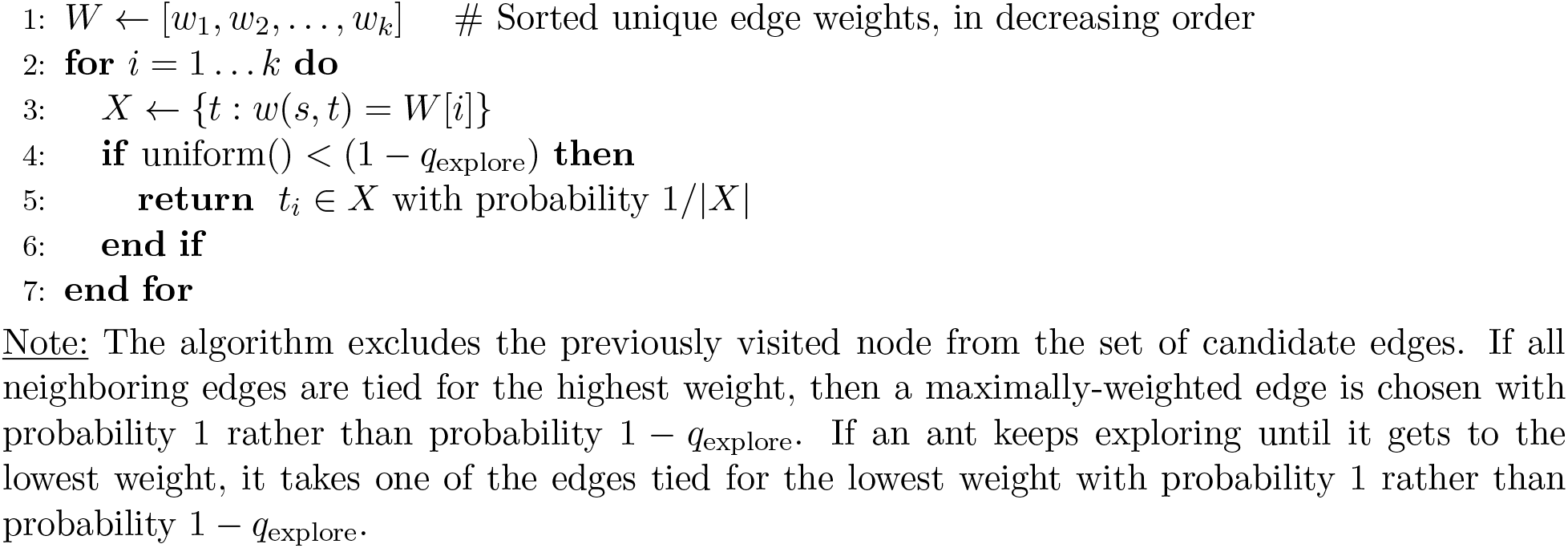
RANKEDGE random walk.

Each algorithm contains additional details inspired by field observations, including a queueing system so ants traverse edges one at a time, the ability to traverse and return from an edge on an explore step in one time-step, and the ability to discontinue laying pheromone on the way back from a dead-end. See Methods for full details.

#### Other algorithms

We compared these two candidate distributed algorithms to several other nonlinear algorithms (MAXEDGEA, MAXEDGEB, MAXEDGEC, MAXWEIGHTED, and DENEUBOURG), as described in the Supplement. We also compared to a null model, called the UNWEIGHTED random walk. The null model uses no parameters; instead, the ants ignore edge weights and choose amongst candidate edges with equal probability.

#### Summary of conclusions

Overall, we find that non-linear models perform the best at simultaneously explaining field observations and providing a mechanism by which turtle ants could solve the network repair problem. While the linear (Weighted) algorithm does perform well at explaining some aspects of field observations, and it repairs breaks with high probability, it also produces a very high path entropy, with poor convergence to a consensus path. This departs strongly from field observations [12] that show that when repairing broken paths, the ants quickly converge to a single path.

In particular, we find that: (A) RANKEDGE outperforms all other non-linear algorithms, except for MAXEDGEA, in the likelihood of explaining observed edge choices by turtle ants (Table 1). The log-likelihoods of RANKEDGE and MAXEDGEA were nearly identical. (B) When parameterized by field data, RANKEDGE outperforms all other non-linear algorithms in success rate (Table 2); and (C) RANKEDGE is equivalent to all other non-linear algorithms in path entropy (Table 3). Compared to the linear WEIGHTED algorithm, RANKEDGE has a lower likelihood of explaining the observed edge choices and a lower success rate. However, RANKEDGE performs much better in path entropy, path length, and maintaining the trail in the absence of a break. We emphasize that the strong success rate of WEIGHTED is because pheromone is left essentially on every edge in the graph. This guarantees high success but very poor convergence to a single path. Field experiments show that turtle ants converge strongly to a single path.

**Table 1:** Maximum likelihood estimates for each algorithm. For each algorithm, we show the values of *q*_explore_ and *q*_decay_ that maximize the likelihood of producing the observed choices made by turtle ants in the field. Standard deviation is computed across 13 junctions, each corresponding to field observations of one junction on a given day. All models are significantly more likely to explain the data than the null model (UNWEIGHTED).

| Algorithm | $q_{\text{explore}}$ | $q_{\text{decay}}$ | log-likelihood |
| --- | --- | --- | --- |
| UNWEIGHTED | N/A | N/A | -744.25 |
| WEIGHTED | $0.05 \pm 0.06$ | $0.01 \pm 0.10$ | -345.05 |
| RANKEDGE | $0.20 \pm 0.04$ | $0.02 \pm 0.08$ | -405.42 |
| MAXEDGEA | $0.20 \pm 0.04$ | $0.02 \pm 0.08$ | -402.63 |
| MAXEDGEB | $0.42 \pm 0.04$ | $0.01 \pm 0.09$ | -424.61 |
| MAXEDGEC | $0.23 \pm 0.01$ | $0.02 \pm 0.09$ | -586.87 |
| MAXWEIGHTED | $0.22 \pm 0.01$ | $0.01 \pm 0.09$ | -568.70 |
| DENEUBOURG | $0.78 \pm 0.10$ | $0.01 \pm 0.03$ | -518.18 |

**Table 2:**
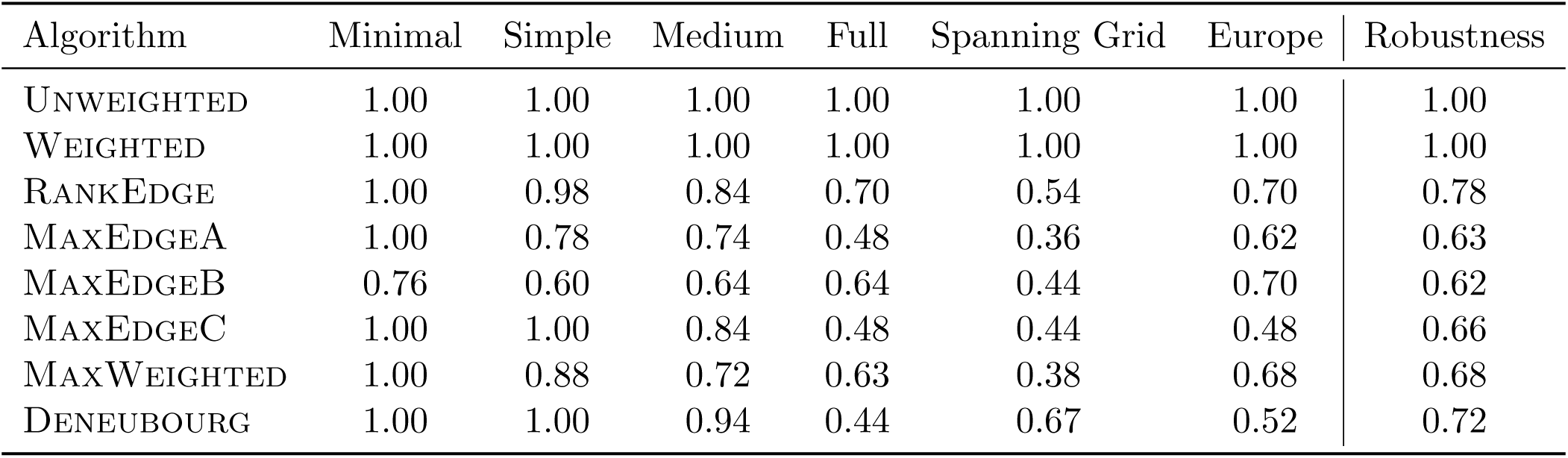
Success rates for each algorithm. For each algorithm, we show the success rate on each simulated network. The last column summarizes the robustness of each method across all networks. Of the non-linear algorithms, RANKEDGE performs the best.

| Algorithm | Minimal | Simple | Medium | Full | Spanning Grid | Europe | Robustness |
| --- | --- | --- | --- | --- | --- | --- | --- |
| UNWEIGHTED | 1.00 | 1.00 | 1.00 | 1.00 | 1.00 | 1.00 | 1.00 |
| WEIGHTED | 1.00 | 1.00 | 1.00 | 1.00 | 1.00 | 1.00 | 1.00 |
| RANKEDGE | 1.00 | 0.98 | 0.84 | 0.70 | 0.54 | 0.70 | 0.78 |
| MAXEDGEA | 1.00 | 0.78 | 0.74 | 0.48 | 0.36 | 0.62 | 0.63 |
| MAXEDGEB | 0.76 | 0.60 | 0.64 | 0.64 | 0.44 | 0.70 | 0.62 |
| MAXEDGEC | 1.00 | 1.00 | 0.84 | 0.48 | 0.44 | 0.48 | 0.66 |
| MAXWEIGHTED | 1.00 | 0.88 | 0.72 | 0.63 | 0.38 | 0.68 | 0.68 |
| DENEUBOURG | 1.00 | 1.00 | 0.94 | 0.44 | 0.67 | 0.52 | 0.72 |

**Table 3:** Path entropy for each algorithm. For each algorithm, we show the path entropy on each simulated network. Lower values indicate convergence to fewer paths. All non-linear algorithms achieve the optimal path entropy. The standard deviation of the entropy for all nonlinear models is 0. We do not report an interval for UNWEIGHTED because it does not depend on pheromone amount and thus has the same limiting behavior in all cases.

| Algorithm | Minimal | Simple | Medium | Full | Spanning Grid |
| --- | --- | --- | --- | --- | --- |
| UNWEIGHTED | 0.00 | 0.69 | 1.79 | 13.75 | 7.24 |
| WEIGHTED | $0.00 \pm 0.00$ | $0.23 \pm 0.25$ | $1.52 \pm 0.20$ | $12.38 \pm 0.22$ | $5.94 \pm 0.15$ |
| RANKEDGE | 0.00 | 0.00 | 0.00 | 0.00 | 0.00 |
| MAXEDGEA | 0.00 | 0.00 | 0.00 | 0.00 | 0.00 |
| MAXEDGEB | 0.00 | 0.00 | 0.00 | 0.00 | 0.00 |
| MAXEDGE C | 0.00 | 0.00 | 0.00 | 0.00 | 0.00 |
| MAXWEIGHTED | 0.00 | 0.00 | 0.00 | 0.00 | 0.00 |
| DENEUBOURG | 0.00 | 0.00 | 0.00 | 0.00 | 0.00 |

## Q1. Field observations to determine the best algorithm and parameter values

We first determined what parameter values best allow each algorithm to match the data from field observations. We then used these parameter values to test algorithm performance for the network repair problem.

The performance of each candidate algorithm is sensitive to the values chosen for the two free parameters, *q*_explore_ and *q*_decay_ (as previously mentioned, we set *q*_add_ = 1). For example, with low values of *q*_explore_, the ants may take a long time to explore enough new edges to find an alternative path; on the other hand, for high values of *q*_explore_, the ants will scatter throughout the network and may not converge to a single path. Similarly, for high values of *q*_decay_ (pheromone decays rapidly), it may be difficult to build and reinforce a single path; for low values of *q*_decay_, it may be hard for the colony to eliminate unnecessary edges and commit to one path. These two parameters also affect each other; for example, the higher the decay rate, the fewer edges with pheromone, and thus the more possible edges to explore.

We used data from observations made in the field to evaluate the match between the choices of edges made by turtle ants and the choices predicted by a candidate algorithm (with parameters *q*_explore_, *q*_decay_). Observations were made at La Estacion Biologica de Chamela in Jalisco, Mexico [6, 12]. Ants were observed traversing a junction (node) along a foraging trail. We recorded the time at which an ant moved to or from that junction node, and the edge it chose to traverse (Figure 2A–B). Observations were made of six different colonies, with an average of 2.16 junctions per colony, over three days in June 2015 and one day in June 2016. We observed 13 different junctions for time periods ranging from 7 to 24 minutes (mean of 12.3 minutes per observation at a given colony on one day), for a total of 773 edge choices made by turtle ants.

### Maximum likelihood estimation

We determined which algorithm and parameter values best explained the observed edge choices made by turtle ants using maximum likelihood estimation (MLE). The data were used to determine the likelihood that a given algorithm, with a given pair of parameter values, would have produced the observed set of edge choices. Figure 2A–B shows an example likelihood calculation, and Figure 2C–D illustrates the results of the MLE for each algorithm over all pairs of parameter values.

Overall, for RANKEDGE, the maximum likelihood parameter values that best explained the observed turtle ant behavior were: *q*_explore_ = 0.20, and *q*_decay_ = 0.02 (Table 1). For WEIGHTED, the maximum likelihood parameter values were *q*_explore_ = 0.05 and *q*_decay_ = 0.01. Both candidate algorithms were more likely to explain the data than the null model (Table 1).

### Consistency of the maximum likelihood estimation across colonies and days

The maximum likelihood parameter values were similar across colonies and days for the 13 junctions (Figure S2). This suggests that across six colonies, there are similar chemical properties in the pheromone (related to *q*_decay_), and that a similar search strategy is used for choosing which edge to traverse next (related to *q*_explore_).

## Q2. Algorithm performance on synthetic and real-world planar networks

Our goal here is to test how well each algorithm solves the network repair problem on simulated and real-world networks. We were particularly interested in how well each algorithm performed when its parameters were set to the maximum likelihood values derived from observations of turtle ants. Our main result is that the maximum likelihood parameters for the RANKEDGE performed well in simulations for network repair across six networks (Figure 3; the black rectangle in both panels shows that the parameter values that best explain the turtle ants’ behavior also perform best for solving the network repair problem.) The latter result is substantiated below.

### Simulation setup

For all simulations, we ran each algorithm for *T* = 1000 steps using *N* = 100 ants, and repeated each simulation 50 times. To initialize each simulation, we placed each of the *N* ants at a random node in the original path. This means that at the start of the simulation there were likely ants at nodes on both sides of the rupture in the path. No ants were placed at nodes not part of the original path. Each ant was randomly assigned to walk in search of one of the two nests. All edges that were part of the initial path were initialized with 10 units of pheromone. All other edges were initialized to 0 units of pheromone. When an ant reached its destination nest, it attempted to return to the other nest, and repeated this, going back and forth between nests, for *T* time-steps. The ants walk synchronously for *T* time-steps; this is a common assumption in distributed computing problems.

Our first performance metric, called the *success rate*, measures how well the ants succeed in finding an alternative path to repair the break. We simulated breaks under six planar network structures, which have an increasingly complex topology with varying numbers of possible paths. In each evaluation below (Figure 4), we show three panels: the initial network with a break, the final network at the end of the simulation, which is generated using the MLE parameter values, and a heatmap showing the success rate for pairs of parameter values (*q*_explore_, *q*_decay_) close to the MLE range. In each synthetic network, a only single link is broken (shown as the ‘X’ mark in Figure 4); in the Supplement, we describe cases where multiple links are broken.

We analyzed all algorithms but show results only for RANKEDGE in the main text because WEIGHTED rarely converged onto a single path, thus it did not satisfy our second performance metric. It also did not maintain trails in the absence of a break. These results are described in detail below. Also, see Table 2 and Supplement for analysis of the additional non-linear algorithms.

### Minimal graph (Figure 4A)

Here we find that RANKEDGE can solve a basic repair problem in a minimal working example, in which the break causes the existing path to lead to a dead end that should be avoided in favor of a single alternative path to the nest. To favor the alternative path, the simulated ants must largely eliminate the pheromone on the edge leading to the dead end, a process which we call ‘pruning’. To favor the alternative path instead of the existing path, the ants should put more pheromone on the edge leading upwards to the alternative route, even though this edge initially had no pheromone.

We find that the RANKEDGE algorithm succeeds in this task 100% of the time, as long as the ants do not leave pheromone on the way back returning from the dead-end (Methods and Q4).

### Simple graph (Figure 4B)

Here we increased the complexity of the graph to offer two alternative paths, instead of one in the Minimal graph. We found that RANKEDGE not only prunes the deadend, but it can find and commit to one of the two alternatives with a 98% success rate.

### Medium graph (Figure 4C)

Here we further increased the complexity of the Minimal graph to offer six alternative paths and found that RANKEDGE not only prunes the dead-end, but can find and commit to one of the six alternatives with a 84% success rate.

### Full grid (Figure 4D)

The Full grid presents a different computational challenge: there is no deadend to prune and the shortest alternative path requires only 3 additional edges. However, the total number of possible new paths is extremely large, which makes it difficult to find and commit to a single path. The Full grid is also a standard benchmark used in the ANTS problem (Related work), in which ants search the integer plane [38, 39, 40].

We found that the highest success rate (70%) occurred for low values of *q*_decay_, which closely matches the observed best decay value estimated using maximum likelihood. This highlights an inherent trade-off in the turtle ant algorithm. Low decay rates help preserve the initial path and bias the turtle ants toward finding an alternative route that re-uses as much of the previous path as possible; that is, with low decay rates, repair starts as close to the break as possible. However, low decay rates also limit the capacity to search for other paths that may be shorter even though they re-use less of the previous path. An alternative would be to use higher values of *q*_explore_ to search for other paths that do not re-use the initial path, but this would make it more difficult for the ants to converge to a single new path.

### Spanning grid (Figure 4E)

In contrast to the Full grid, the Spanning grid is sparser and requires that the ants go back at least one node from the break to find an alternative path.

We found that the maximum likelihood parameters produced a moderate success rate (54%). As above, the highest success rate occurred for low values of *q*_decay_ and moderate values of *q*_explore_. These values achieve a good trade-off between searching sufficiently far from the break to find an alternative path, and largely preserving the previous path. The performance on the Spanning grid demonstrates that the algorithm is flexible enough to search locally around a break point for new paths, while still maintaining most of the old path.

The results from the Full grid and the Spanning grid together suggest that the algorithm performs best when it preserves as much of the previous path as possible, even if it can not re-use all of the original path. This is consistent with field observations that showed that turtle ants sought alternative paths in a “greedy” manner, by going back up to 1 or 2 nodes from the break point, even though going back more nodes may have resulted in a path with fewer nodes overall [12].

### European road transportation network (Figure 5)

To demonstrate the utility of this algorithm in a real-world scenario, we applied the RANKEDGE algorithm to repair networks in a human-designed transport network. We downloaded the network depicting the major roads (edges) connecting intersections (nodes) in the international E-Road in Europe [62] (Methods). We removed an edge from an existing path between two nodes and ran the RANKEDGE algorithm to repair the simulated closure. The RANKEDGE algorithm achieved a success rate of 70%, indicating that the turtle ant algorithm can also repair breaks in real-world topologies. This shows how distributed solutions may be useful for new application domains, such as for swarm robotics or molecular robots [63, 64, 65, 66] in remote environments, when centralized or global positioning systems may not be as effective.

We also compared the algorithms on Erdos-Renyi random networks and small-world networks and found similar gains in performance for RANKEDGE (Supplement).

### Q2a. Converging onto a single consensus path (Table 3)

Our second performance metric, called *path entropy*, measures how well foragers commit to a single alternative path.

We find that the RANKEDGE algorithm consistently achieves a path entropy near 0, indicating that on the final network all ants follow the same path when not taking explore steps (Table 3). This is particularly challenging for the Full and Spanning grids because both contain a large number of possible paths, and thus a large possible path entropy if the simulated ants exploit many paths. Thus, when the algorithm succeeds in repairing the path, RANKEDGE satisfies our second performance criterion.

More generally, one advantage of non-linear algorithms such as RANKEDGE is that all simulated ants, by simply following the maximal edge (the adjacent edge with the highest pheromone), can travel from one nest to the other using the same path, thereby achieving a path entropy of 0. On the other hand, linear algorithms such as WEIGHTED do succeed in finding a path; however, WEIGHTED is unable to commit to only one path, and thus has very high path entropy (Figure 6).

### Q2b. Finding short paths (Table 4)

Our third performance metric measures the path length of the final trail network. We found that RANKEDGE consistently finds paths of lengths that are close to, though slightly larger than, the globally shortest path lengths. For every network, we compared the average path lengths of RANKEDGE versus every other algorithm using Welch’s unpaired T-test. RANKEDGE finds significantly shorter paths than WEIGHTED and UNWEIGHTED on the Full grid, Spanning grid, and European roads (*p <* 0.05); RANKEDGE is not significantly different from the other non-linear algorithms (Table 4). The improved performance over UNWEIGHTED demonstrates the value of using pheromone to solve the network repair problem collectively, instead of using independent search.

**Table 4:** Average path length for each algorithm. For each algorithm, we show the average path length of the final graph, measured as the number of nodes in the path. RANKEDGE performed much better than the null model (UNWEIGHTED) and close to the globally shortest path length (Optimal).

| Algorithm | Minimal | Simple | Medium | Full Grid | Spanning Grid |
| --- | --- | --- | --- | --- | --- |
| UNWEIGHTED | $12.00 \pm 0.00$ | $12.90 \pm 1.00$ | $13.16 \pm 3.00$ | $26.04 \pm 6.56$ | $20.05 \pm 7.24$ |
| WEIGHTED | $12.00 \pm 0.00$ | $12.97 \pm 0.85$ | $13.04 \pm 0.64$ | $18.36 \pm 0.18$ | $16.93 \pm 0.24$ |
| RANKEDGE | $12.00 \pm 0.00$ | $12.98 \pm 1.01$ | $10.95 \pm 1.01$ | $13.06 \pm 0.34$ | $14.56 \pm 3.97$ |
| MAXEDGEA | $12.00 \pm 0.00$ | $12.87 \pm 1.00$ | $11.29 \pm 0.97$ | $13.17 \pm 0.56$ | $15.56 \pm 4.38$ |
| MAXEDGEB | $12.00 \pm 0.00$ | $13.30 \pm 0.96$ | $10.77 \pm 0.99$ | $13.06 \pm 0.35$ | $15.09 \pm 4.07$ |
| MAXEDGE C | $12.00 \pm 0.00$ | $13.02 \pm 1.01$ | $11.00 \pm 1.01$ | $13.16 \pm 0.56$ | $15.45 \pm 3.54$ |
| MAXWEIGHTED | $12.00 \pm 0.00$ | $12.87 \pm 0.99$ | $10.89 \pm 1.03$ | $13.06 \pm 0.58$ | $14.46 \pm 3.07$ |
| DENEUBOURG | $12.00 \pm 0.00$ | $13.16 \pm 1.00$ | $11.06 \pm 1.30$ | $14.18 \pm 3.47$ | $13.94 \pm 2.51$ |
| Optimal | 12.00 | 12.00 | 10.00 | 13.00 | 13.00 |

### Q2c. The power of bi-directional search (Figure S3)

We find that a bi-directional search, in which simulated ants attempt to create an alternative path concurrently from both sides of the break, allows the algorithm to perform significantly better than a uni-directional search using ants from only one side of the break. We tested this on the Full grid, and found that for the MLE parameter values for RANKEDGE, the success rate was on average 70% for a bi-directional search versus 14% for uni-directional search.

One might predict that uni-directional search would perform as well as the bi-directional search, while simply taking longer. However, we found this not to be true: using a bi-directional search means that once ants from side A of the break reach side B of the break, the rest of their search is directed by the pheromone trail laid by ants that started on side B. In the uni-directional search, even if ants from side A reach side B, they must still find a path from scratch connecting the dead end on side B to the nest on side B. Although uni-directional search has rarely been observed to occur in turtle ant networks, we tested it here to compare it with bi-directional search, which is often used to improve the performance of search algorithms.

### Q2d. The power of avoiding backtracking (Figure S4)

We find that providing simulated ants the ability to avoid backtracking, i.e., visiting the same node visited in the previous time-step (Methods) allows for a significant improvement in algorithm performance. In contrast, ants that are not given this ability could keep going back and forth along the same edge.

In particular, ants that used the RANKEDGE algorithm and avoided backtracking produced a success rate of 70% on the Full grid, compared to 0% when an ant was not prevented from potentially returning to the previous node it visited (Figure S4). Thus, providing ants with a basic node-to-node sense of direction led to a significant improvement in performance.

### Q2e. Critical features for the success of a plausible algorithm

We find that there are two important features for non-linear algorithms to perform well in simulation: (1) Simulated ants do not lay pheromone on the way back from a dead end, and (2) simulated ants queue at nodes (Methods). In the Supplement, we provide theoretical analysis for why these two components are critical for any non-linear algorithm to circumvent a dead end. Without either of these two features, the time to circumvent a dead end rises dramatically. In the Supplement, we also confirm these theoretical observations via simulation.

## Q3. Maintaining a trail in the absence of a break

Here we consider whether the same algorithm used to repair a path can also keep a path intact when it is not broken. This is important because if different algorithms were used to maintain trails versus repair trails, then the turtle ants would need some signal to toggle between different methods for choosing among candidate edges, depending on the context. We found that a single algorithm, RANKEDGE, is capable of maintaining trails and responding to breaks.

In particular, we ran the RANKEDGE algorithm on the Spanning grid without breaking the original path and found that the trail was preserved without any modification to the algorithm or its parameters (Figure 7). In contrast, the WEIGHTED algorithm performed very poorly on this task. In particular, for RANKEDGE, the path entropy using the MLE parameter values from turtle ant data was optimal (0.00). For WEIGHTED, however, the path entropy for the MLE parameter values was much higher (5.38), indicating poor maintenance of the original path.

## Q4. Pruning as a general principle for discovering alternative paths

Field observations show that turtle ants engage in pruning (Figure 8). In our simulations, we also observed that ants explored multiple alternative paths, and then most of these paths were pruned as the colony converged to a single alternative path. Further, the paths tended to become shorter over time. We quantified how many paths were pruned during the simulation using a measure called *path elimination* (Methods). We also quantified how the lengths of the remaining paths changed over time using a measure called *path length pruning* (Methods). All of the non-linear algorithms exhibit some path elimination and pruning, and thus could plausibly be used to explain the observed pruning in observed turtle ants. RANKEDGE prunes fewer paths than the other algorithms (Table 5) because RANKEDGE does not form as many initial paths as other algorithms. However, of the paths that are pruned, RANKEDGE tends to prune more nodes from the paths (Table 6).

**Table 5:**
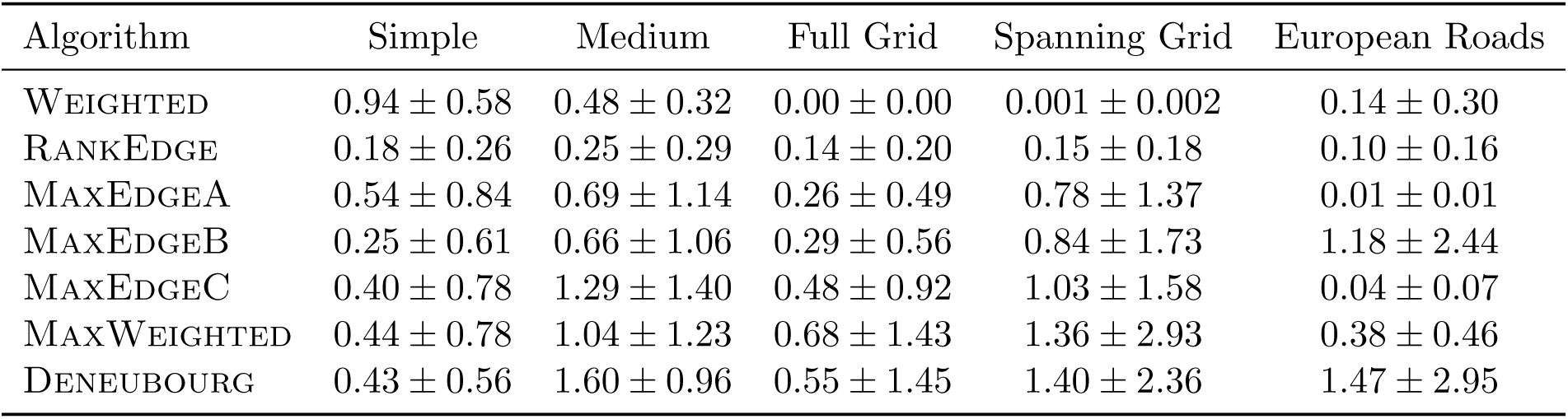
Path elimination: For each algorithm, we show the average reduction in entropy over chosen paths over time (Methods). We omit the Minimal graph, because there is only one possible path from one nest to the other, and thus no path elimination is possible.

| Algorithm | Simple | Medium | Full Grid | Spanning Grid | European Roads |
| --- | --- | --- | --- | --- | --- |
| WEIGHTED | $0.94 \pm 0.58$ | $0.48 \pm 0.32$ | $0.00 \pm 0.00$ | $0.001 \pm 0.002$ | $0.14 \pm 0.30$ |
| RANKEDGE | $0.18 \pm 0.26$ | $0.25 \pm 0.29$ | $0.14 \pm 0.20$ | $0.15 \pm 0.18$ | $0.10 \pm 0.16$ |
| MAXEDGEA | $0.54 \pm 0.84$ | $0.69 \pm 1.14$ | $0.26 \pm 0.49$ | $0.78 \pm 1.37$ | $0.01 \pm 0.01$ |
| MAXEDGEB | $0.25 \pm 0.61$ | $0.66 \pm 1.06$ | $0.29 \pm 0.56$ | $0.84 \pm 1.73$ | $1.18 \pm 2.44$ |
| MAXEDGE C | $0.40 \pm 0.78$ | $1.29 \pm 1.40$ | $0.48 \pm 0.92$ | $1.03 \pm 1.58$ | $0.04 \pm 0.07$ |
| MAXWEIGHTED | $0.44 \pm 0.78$ | $1.04 \pm 1.23$ | $0.68 \pm 1.43$ | $1.36 \pm 2.93$ | $0.38 \pm 0.46$ |
| DENEUBOURG | $0.43 \pm 0.56$ | $1.60 \pm 0.96$ | $0.55 \pm 1.45$ | $1.40 \pm 2.36$ | $1.47 \pm 2.95$ |

**Table 6:** Path length pruning: For each algorithm we show the average reduction in lengths of chosen paths over time (Methods) observed. We omit the Minimal graph, because there is only one possible path from one nest to the other, and thus no pruning is possible.

| Algorithm | Simple | Medium | Full Grid | Spanning Grid | European Roads |
| --- | --- | --- | --- | --- | --- |
| WEIGHTED | $0.94 \pm 0.58$ | $0.48 \pm 0.32$ | $0.00 \pm 0.00$ | $0.001 \pm 0.002$ | $0.14 \pm 0.30$ |
| RANKEDGE | $0.30 \pm 0.58$ | $0.39 \pm 0.58$ | $0.04 \pm 0.14$ | $0.97 \pm 2.59$ | $0.35 \pm 0.50$ |
| MAXEDGEA | $0.30 \pm 0.27$ | $0.28 \pm 0.30$ | $0.11 \pm 0.17$ | $0.15 \pm 0.20$ | $0.05 \pm 0.10$ |
| MAXEDGEB | $0.14 \pm 0.23$ | $0.15 \pm 0.23$ | $0.06 \pm 0.10$ | $0.11 \pm 0.16$ | $0.08 \pm 0.14$ |
| MAXEDGE C | $0.31 \pm 0.29$ | $0.50 \pm 0.36$ | $0.12 \pm 0.17$ | $0.13 \pm 0.21$ | $0.08 \pm 0.13$ |
| MAXWEIGHTED | $0.29 \pm 0.30$ | $0.39 \pm 0.33$ | $0.15 \pm 0.23$ | $0.19 \pm 0.24$ | $0.17 \pm 0.23$ |
| DENEUBOURG | $0.46 \pm 0.17$ | $0.42 \pm 0.28$ | $0.001 \pm 0.003$ | $0.04 \pm 0.09$ | $0.02 \pm 0.09$ |

For every network, we compared the average path elimination and path length pruning of RANKEDGE versus every other algorithm using Welch’s unpaired T-test. For path elimination, RANKEDGE does not reduce the number of paths more than other algorithms (*p <* 0.05), as we described above. However, for path length pruning, RANKEDGE reduces path lengths significantly more than all other algorithms on the Spanning grid and European roads (*p <* 0.05). On the Full grid, RANKEDGE is significantly better than WEIGHTED (*p <* 0.05), is not significantly different from DENEUBOURG or MAXEDGEB, and is significantly worse than MAXEDGEA, MAXEDGEC, and MAXWEIGHTED. No statistical difference is observed for the Simple and Medium graphs, likely due to their relatively simple topology. These pruning results suggest that RANKEDGE explores fewer paths initially but is better at selecting the shortest of the paths it explores. RANKEDGE tends to prune fewer paths (path elimination), but of the paths it does explore, RANKEDGE converges to the shorter paths (path length reduction).

Field observations [12] also showed that when ruptured trails are repaired, new nodes are added to the network, and in subsequent days, some of the nodes are pruned. Such pruning of nodes also led to global pruning of paths (Figure 8). Such an “explore-exploit” strategy may help turtle ants quickly find a solution that re-connects a rupture in a trail, and may also help the colony to optimize the coherence of the trail, by minimizing the number of junctions at which ants could get lost.

Interestingly, using pruning-based strategies to discover the most appropriate edges or paths to keep is a common strategy used by biological systems. In particular, during the development of neural circuits in the brain, synapses are massively over-produced and then pruned-back over time [3]. This strategy is thought to help neural circuits explore possibly topologies and then converge to the most appropriate topology based on environment-dependent feedback. A similar process occurs during the development of vascular (blood flow) networks in the body [67]. Thus, pruning may be a common biological strategy of network design when multiple topologies need to be explored in a distributed manner.

## Discussion

Our primary contribution is to address an engineering problem (maintaining a trail network and finding alternative paths to route around broken links in the network) using biologically feasible parameters and models motivated by how turtle ants may solve this problem in the field. Successful performance by the algorithm in simulation, using realistic parameter values, indicates that the algorithm is a plausible candidate to describe how the ants create their networks. The RANKEDGE algorithm achieved a better maximum likelihood estimate (Figures 2–3) than every other non-linear model except for MAXEDGEA. When parameterized by data from field observations, RANKEDGE was better able to find a single, short path with high probability compared to other algorithms (Tables 2,3, 4). From this, we conclude that non-linear models, in particular RANKEDGE and MAXEDGEA, represent the two best plausible models of turtle ant behavior.

By testing performance across six different networks, we found that the turtle ants appear to have evolved an algorithm that may not be optimal for any particular planar network but is robust to some variation in the topology. Further, non-linear algorithms exhibited pruning, which also occurred in field observations [12] (Table 6, Figure 8).

There are several features of our algorithm that are critical for success in repairing breaks to the routing backbone. First, to minimize path entropy it is essential to have a stronger-than-linear bias towards choosing the highest-weighted edge (RANKEDGE), rather than choosing edges proportional to their edge weight (WEIGHTED). This helps constrain the search space and leads to better convergence to a single consensus path. We emphasize that WEIGHTED has a strong success rate because pheromone is left on every edge in the graph (see e.g., Figure 6A–B). This guarantees that there exists some path with positive probability. However, WEIGHTED does not commit to a single path as effectively as an algorithm with a strong, non-linear bias toward the highest-weighted edge, such as RANKEDGE. The path entropy of WEIGHTED is high because essentially every path in the graph has high probability, and the average path length is high because the algorithm does little to eliminate long paths and commit to short paths. Second, to repair breaks it is essential to use bi-directional search and avoid backtracking. Observations show that when turtle ants encounter a break, ants from both sides of the break attempt to repair the trail [12]. When ants from both sides meet, they each encounter a trail that is already strongly reinforced and guided towards the other nest. In addition, the ability to avoid backtracking allows ants to avoid going back and forth along the same edge. Third, we showed theoretically that the time needed to find an alternative path decreases significantly if turtle ants reaching a dead-end in their trail do not leave pheromone while returning back from the dead-end.

The RANKEDGE algorithm is parsimonious, capable of both maintaining trails and repairing breaks to trails using the same underlying logic. Observed ants encounter diverse situations analogous to breaks in the ongoing maintenance of trails. We find that a single algorithm can solve two diverse problems without requiring the additional complexity of a signal that distinguishes such situations from a rupture in the trail. How each path is established originally is an interesting yet distinct question. Paths are not always the shortest globally, and the physical structure of edges in the canopy appears to affect how these paths are selected.

The algorithm can be extended to improve performance, though this may involve sacrificing biological realism. One possible extension would allow ants to “toggle” between different parameter values or algorithms in different situations. For example, an ant could use RANKEDGE, but if it encounters a dead-end or massive crowding (determined for example by a large increase in the frequency of antennal contacts with other ants [68, 18]), then it increases its probability of exploring new edges. This would be similar to a distributed version of simulated annealing, with the value of *q*_explore_ corresponding to the decreasing value of the temperature parameter. A second possible extension would be to use multiple types of pheromone [69]. Ants could use negative pheromone to signal to other ants not to select a certain edge, for example, towards a dead-end. Further work is needed to measure the computational abilities of turtle ants to determine whether such extensions depart from biological realism.

There are some differences between our synthetic networks and the environment of the observed turtle ants. First, turtle ant trail networks in the canopy are 3D planar networks, whereas here, to begin the investigation of arboreal ant trail networks, we used 2D planar networks. The ideal test case would be a suite of synthetic networks that are isomorphic to some portion of the turtle ant canopy. In lieu of this, we describe five synthetic networks, some of which have been used by prior work, that each collectively test the ability of different algorithms to solve the network repair problem. These five networks comprise a necessary (if not exhaustive) set of test cases. Second, in the tropical forest, many edges are physically difficult to traverse, which may provide natural inhibition for selecting certain edges. Third, in the canopy, edges are not all of the same length. In future work that includes variability in edge lengths, synchronous walks will need to be modified since longer edges require more time-steps to traverse. More generally, further work is needed to determine the physical properties of junctions and branches in the canopy and how these properties influence the likelihood of traversing an edge.

Finally, the probabilistic RANKEDGE algorithm is biologically feasible, requiring less computational complexity and assuming fewer memory requirements than many other distributed graph algorithms commonly used in computer science. This suggests that a biological algorithm evolved to deal with the constraints of the tropical forest canopy may be useful in other applications, such as in swarm robotics or molecular robots [63, 64, 65, 66]. For such applications, the best algorithm to choose depends on the requirements of the problem. We find evidence that RANKEDGE is the best algorithm to achieve relatively high success rate and low path entropy. If agents do not need to adhere to a single path, then the WEIGHTED algorithm performs better, though the length of the path may be long. It appears that turtle ants use an algorithm that finds a single short path [12], as RANKEDGE provides. For trivial graphs with only one alternative path, both algorithms perform similarly.

Overall, our work contributes to the growing intersection of distributed algorithms used by natural biological processes [70, 11].

## Methods

### Maximum likelihood estimation of parameter values

For each candidate algorithm, we varied *q*_decay_, *q*_explore_ *∈* (0, 1) and evaluated the likelihood that the algorithm with a specific set of parameter values would have generated the choices made by turtle ants observed in the field. The edges traversed by the turtle ants, and times the edges were traversed, were used to compute how much pheromone had been added to and had decayed from each edge, to give the amount of pheromone on each edge at any time. In modeling pheromone decay, we treat *q*_decay_ as the rate of decay per second. When computing the amount of decay between two consecutive ant choices, we decay all of the edges in proportion to the number of seconds elapsed between the two choices. For each candidate algorithm, if we know all of the edge weights at a given time and the value of *q*_explore_, we can compute the likelihood of a given choice. Figure 2A–B provides an example of a calculation of the likelihood of a choice for each candidate algorithm.

For a given combination of *q*_decay_, *q*_explore_ we performed this likelihood computation for every observed choice in each of the 13 junctions. We updated the edge weights based on the choice and the amount of time that passed between successive choices, and then repeated this process on the next choice made by the next ant. For each junction of observations at a given node on a given day, we computed the maximum likelihood estimate (MLE) for each parameter value pair. We then added the log-likelihoods for all the 13 junctions.

As we formalize in the Supplement, the exponential rate of pheromone decay means that the most recent ant choices have the largest effect on the current edge weights at a junction. Pheromone added far in the past will have largely decayed and will not contribute much to the current weights. Thus, we do not need an extensive history of the choices to perform accurate modeling. It is not currently possible to measure or manipulate pheromone levels on the branches in the canopy.

### Additional technical details

Each algorithm includes the following constraints motivated by field observations:

1. Observations suggest that turtle ants tend not to backtrack, but instead tend to keep moving along the trail in the same direction, indicating that turtle ants have at least enough sense of direction to avoid going back and forth over the same edge. Our simulations include three exceptions to this. First, because our simulations include two nests with ants going back and forth, upon reaching the nest, an ant is allowed to backtrack along the same edge it used to reach the nest. Second, if a simulated ant reaches a dead-end node that has no outgoing edges other than the previously traversed edge, it is allowed to backtrack. However, the ant does not lay pheromone on the way back until it reaches a node with two edges, excluding the edge it previously traversed. In field experiments, it is difficult to determine whether a turtle ant is laying pheromone; however, it is known that *Lasius niger* ants down-regulate pheromone deposition at dead-ends to avoid recruitment during crowding [71, 69, 69]. It is possible that turtle ants similarly down-regulate pheromone in response to dead-ends. In the Supplement, we also provide a probabilistic argument for why it is critical that ants do not lay pheromone when returning from a dead-end to repair the break. Thus, in addition to the ability to avoid backtracking, each ant requires one binary state variable that is 0 or 1 depending on whether the ant is coming back from a dead end.
2. Turtle ants queue at a node and leave in a first-in first-out manner. In other words, if more than one ant is at the same node, only one ant chooses an edge in each time-step. In the field, turtle ants walk along narrow branches, almost always one ant at a time in each direction. We find that queueing increases the success rate of the algorithm (Supplement).
3. When turtle ants take an “explore step”, they often traverse an edge for a short distance, and then return to the original node [12]. This builds a slight extension off the primary path, which can be extended by subsequent ants. In all algorithms, if a simulated ant takes an explore step, it goes across the edge and comes back in one time-step. Thus, two units of pheromone are left on the edge, and the ant is back at the node it started from. (R3-27) An explore step is defined as any choice that cannot occur unless *q*_explore_ *>* 0. For WEIGHTED, this involves taking an edge with zero weight; for RANKEDGE, this involves taking an edge that does not have the highest weight. For UNWEIGHTED, there is no explore step, because ants are not inherently biased towards following any particular pheromone trail.
4. Because pheromone decays exponentially, theoretically once an ant lays pheromone on an edge, that edge’s weight will never decay to absolute 0. In practice, if the edge weight stays unchanged even after multiplying by the decay rate (due to numerical computation error, occurring at roughly 10^*−300*^), then we reset the weight of that edge to absolute 0. Another possible approach would be to introduce a pheromone detection threshold parameter. If an edge had pheromone below this threshold, the ant would treat the edge as if it had no pheromone. We avoided this approach because it would introduce another parameter to optimize and compare.
5. All edges are assumed to have the same length, and it takes exactly one time step for an ant to cross an edge.

### Performance metrics

Below we formally describe the three performance metrics used to evaluate each algorithm, after it ran for *T* time-steps. Intuitively, the simulated ants have successfully found a path if they can reach one nest from the other without taking any “explore steps”. We thus measure the performance of each algorithm assuming *q*_explore_ = 0, and consider all paths that may be taken with positive probability under this constraint. For RANKEDGE and other non-linear algorithms, ants travel from one nest to the other by following the highest-weighted edges. For WEIGHTED, ants travel from one nest to the other using only edges of positive weight.

Formally, let the *pheromone subgraph* be the subgraph induced by the two nest nodes, all edges with a nonzero weight, and all nodes adjacent to edges with non-zero weight. Let *G* be a pheromone subgraph and *P* = (*v*_1_, *v*_2_, *… v*_*n*_) be a path in *G*, with nests *v*_1_ and *v*_*n*_. For a node *v*_*i≠*1_ *∈ P*, define the candidate edges *C*_*P*_ (*v*_*i*_) = *{*(*v*_*i*_, *u*) *∈ E*(*G*) : *u ≠ v*_*i-*1_*}*, i.e., the edges that the ant could take from *v*_*i*_ without backtracking to its previous node. Let *v*_*i*_, *v*_*i*+1_ *∈ P* be consecutive nodes in the path; we say edge (*v*_*i*_, *v*_*i*+1_) is *maximal with respect to P* if *w*(*v*_*i*_, *v*_*i*+1_) = max*u∈CP* (*vi*) *w*(*v*_*i*_, *u*). The path *P* is a *maximal path* if for every pair of consecutive nodes *v*_*i*_, *v*_*i*+1_ *∈ P*, the edge (*v*_*i*_, *v*_*i*+1_) is maximal with respect to *P*. An ant taking a maximal path always takes an edge with the highest weight; thus, a maximal path allows an ant using one of the non-linear algorithms to commute between two nests with positive probability even if *q*_explore_ = 0.

Next, define a *pheromone path* to be a path in which all edges have positive weight. Such a path allows an ant following the WEIGHTED algorithm to commute between two nests with positive probability even when *q*_explore_ = 0.

For a given algorithm, define a *solution path* to be a path that can be traversed with positive probability under that algorithm when *q*_explore_ = 0. For all the non-linear algorithms discussed here, solution paths are equivalent to maximal paths. For the WEIGHTED algorithm, a solution path is equivalent to a pheromone path.

Let 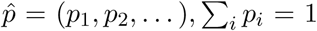 be a probability distribution. Define the *entropy* of the distribution to be: *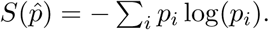*

At the end of the simulation, we evaluate the pheromone subgraph of each algorithm by computing the following measures:

- **Success rate (higher is better):** The probability that the ants form a solution path. This is defined empirically by computing the percentage of the *N* simulations where a solution path is formed in the final graph.
- **Path entropy (lower is better):** An information-theoretic measure of how well the ants converge onto a single solution path.
  - Let *M*_1_, *M*_2_, *…, M*_*n*_ be the set of all *n* solution paths in the final graph.
  - Let *p*_1_, *p*_2_, *…, p*_*n*_ be the probabilities of taking each solution path with *q*_explore_ = 0. The probabilities 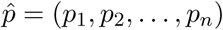 form a probability distribution.
  - The path entropy is then: *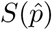*
- **Average path length (lower is better):** The average length of the solution paths in the final graph. Path length is defined to be the number of nodes in a path.

To compute the pruning metrics, we first define a *chosen path* as the sequence of nodes *v*_1_, *v*_2_, *…, v*_*n*_, after removing cycles, that an ant takes to successfully walk from one nest to another. Figure S1 illustrates why removing cycles is necessary when comparing chosen paths.

Over the course of the simulation, we track all chosen paths for all ants that successfully walk from one nest to the other. This includes the number of times each path was chosen — and thus the distribution over the chosen paths — and the lengths of these paths.

- **Path elimination:** An information-theoretic measure of the degree to which paths from one nest to the other are eliminated over time.
  - Let *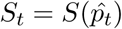* be the entropy over the distribution of chosen paths 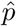 that have been completed at or before time *t*.
  - Let 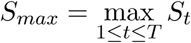 be the maximum chosen-path entropy over the entire simulation.
  - The path elimination is then the maximum entropy minus the entropy at the end of the simulation: *S*_*max*_*− S*_*T*_.
- **Path length pruning**: A measure of the degree to which ants reduce the lengths of the paths they take over time.
  - Suppose at time *t* the ants have taken chosen paths *p*_1_, *p*_2_, *…, p*_*n*_ with frequencies *c*_1_, *c*_2_, *…, c*_*n*_. Let *l*(*p*_*i*_) be the length of path *p*_*i*_. We define the *weighted-mean chosen path length* at time *t* to be: *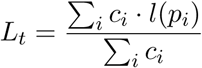.*
  - Let 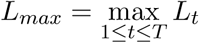 be the maximum weighted mean chosen path length over the entire simulation. simulation.
  - The path length pruning is then: *L*_*max*_ *− L*_*T*_.

#### Robustness across network topologies

To determine which parameter values performed well across all the planar topologies tested, we defined the *robustness* of a set of parameter values (*q*_explore_, *q*_decay_) to be the geometric mean of the success rates for those parameter values on all six networks. We use the geometric mean because it penalizes parameter values that perform poorly on any one particular graph; for a set of parameter values to have a high geometric mean, it must perform well on every graph. When computing robustness, we weight the success rates over all networks equally. This is done for two reasons: first, this highlights algorithms that perform well under a variety of distinct but equal conditions; and second, we do not currently have complete data on which topologies are more or less likely to occur in the canopy, and thus it is not clear how weighting factors should be selected.

#### Application to the European road network

We sampled a portion of the European road network. This sample contained the same number of nodes as the Full grid (11 *×;* 11 = 121 nodes). Sampling was done by selecting a random node and performing a breadth-first search until 121 nodes were visited. The network contained these 121 nodes and all the edges adjacent to these nodes. We then randomly selected two nodes and removed a randomly-chosen edge in the shortest path between those nodes. If removing this edge disconnected the two nodes, we discarded the pair of nodes and picked a new randomly chosen pair of nodes. We then applied our algorithm to repair the trail.

## Figure Legends

Figure 1: **Turtle ant habitat and trail network.** A) The photograph shows the highly tangled forest canopy in which turtle ants forage. B) Experiments were performed in which an edge in the path was cut, to observe how the ants respond and repair the break [12]. C) Modeling the trail network as a graph, with junctions as nodes and connecting branches and twigs as edges. The diagram on the right from [12] shows a detailed depiction of a large portion of the trail network. Each days path is shown in a different color (see legend), and additional repair paths are shown in a distinct color. Solid lines connect two nodes that are on the same plant (e.g. node 36 and node A are on the same plant). Dashed lines connect two nodes that are on a different plant (e.g. nodes B and C are on different plants).

Figure 2: **Maximum likelihood computation.** A–B) Example node junction and edge choices for turtle ants. All ants arrive at node 1 from a different node that is not shown. In the example, we assume pheromone has been deposited at previous time-points, and we now compute the likelihood of the next ant choice. Under the <u>RANKEDGE</u> algorithm, the likelihood of choosing edges 1 *→* 3 or 1 *→* 2 is (1*− q*_explore_)(1*/*2); the likelihood of edge 1 *→* 4 is *q*_explore_(1 *− q*_explore_); and the likelihood of 1 *→* 5 is 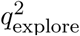. Under the <u>WEIGHTED</u> algorithm, the likelihood of choosing edge 1 *→* 5 is *q*_explore_; the likelihood of edge 1 *→* 4 is (1 *− q*_explore_)(1*/*(1 + 2 + 2)); and the likelihood of edges 1 *→* 2 or 1 *→* 3 is (1 *− q*_explore_)(2*/*(1 + 2 + 2). Under the <u>UNWEIGHTED</u> algorithm, the edge weights are disregarded, and the likelihood of taking any one of the four edges is (1*/*4). C–D) For each combination of *q*_explore_ (*x*-axis) and *q*_decay_ (*y*-axis) values, we determined the pair’s likelihood of producing the choices observed in turtle ants. Each heatmap shows the likelihood for each algorithm with a zoom-in below around the highest likelihood region. The optimal parameter values for each algorithm, depicted in white, are shown in Table 1.

Figure 3: **The maximum likelihood parameters closely match the best simulation parameters:** A) The color of each square in the heatmap corresponds to the robustness (Methods) of the success rates for the RANKEDGE algorithm for each combination of *q*_explore_ (*x*-axis) and *q*_decay_ (*y*-axis) values. Results are aggregated over the six simulated and real-world networks presented in Figures 4 and 5. B) The maximum likelihood parameter estimates for RANKEDGE from observations of turtle ants. The black rectangle in both panels shows that the parameter values that best explain the turtle ants’ behavior also perform best for solving the network repair problem.

Figure 4: **Success rates for each network.** A–E) For each network we show the initial graph (left), an example of the final graph after running the RANKEDGE algorithm using the maximum likelihood parameters (middle), and the algorithm’s success rate for each parameter combination (right). In each panel, black dots indicate nodes in the network, and solid lines indicate edges that may be traversed. If two adjacent nodes are not connected by an edge, there is a space between them. In the initial graphs, the ‘X’ marks the edge that is broken. The *x*-axis of the heatmap (right column) shows *q*_explore_, and the *y*-axis shows *q*_decay_ under the range close to the MLE parameters. Darker shades of red are indicate success rates closer to 1, and thus are better.

Figure 5: **Repairing road closures in the Europe road graph.** Analysis of how well the turtle ant algorithm translates to repair simulated breaks in a real-world transport network. A) An example of a path in the European E-road network connecting Munich to Berlin, Germany. The roads and junctions form a graph. On the left, the black ‘X’ shows a road that has been broken or closed along the path. On the right, we show an alternative path that avoids the broken road. B) The success rate of the turtle ant algorithm (RANKEDGE) applied to this network. Map data: Google, DigitalGlobe.

Figure 6: **Poor path entropy for WEIGHTED.** The initial (left) and final (right) networks for the (A) Full grid and (B) Spanning grid. In both cases, the MLE parameter values (*q*_explore_ = 0.05, *q*_decay_ = 0.01) for WEIGHTED did not find a low path entropy solution.

Figure 7: **Analysis in the absence of a break.** A) Initial Spanning grid, with no break. B) The final network produced using WEIGHTED, which does not find a low entropy solution. C) The final graph using RANKEDGE, which finds a low path entropy solution.

Figure 8: **Turtle ants prune paths.** The diagram from [12] shows the results of an experiment in which an edge was cut. Left: The initial trail is shown in grey. The edge connecting nodes 5 and 6 was cut. After 75 minutes, the turtle ants explored several new paths (red). Center: Five hours after the cut, some of the red paths were pruned (transparent grey). Ants traveling down from node 7 took one trail, consisting of nodes 6, 15, 16, 17, 18, 4, and 3, because they could not use 12 in this direction. Ants traveling in the other direction took another trail, consisting of nodes 3, 4, 5, 12, 13, and 14, or the trail consisting of 11, 13, and 14. Right: The next day, there was additional pruning. Because node 12 could now be used in both directions, ants traveled both ways on the indicated trail.

## Acknowledgments

The authors thank Jason Schweinsberg for guiding us in the theoretical proof. We are grateful to Illia Ziamtsov, Javier How, Benjamin Cosman, Ailie Fraser, Will Hamilton, Sam Crow, Alex Lang, and the anonymous reviewers for helpful comments on the manuscript.

## Source code and datasets

All source code for the algorithm and all datasets for the ant choices are available at our Github repository: http://github.com/arjunc12/Ants.

## Author Contributions

AC and SN performed the computational experiments. DMG performed the field experiments. All authors wrote and reviewed the manuscript.

